# Assessment of machine-learning predictions for MED25 ACID domain interactions with transactivation domains

**DOI:** 10.1101/2023.11.30.569364

**Authors:** Didier Monté, Zoé Lens, Frédérique Dewitte, Vincent Villeret, Alexis Verger

## Abstract

Human Mediator complex subunit MED25 binds transactivation domains (TADs) present in various cellular and viral proteins using two binding interfaces found on opposite sides of its ACID domain, and referenced as H1 and H2. Here, we use and compare deep learning methods to characterize Human MED25-TADs interfaces and assess the predicted models to published experimental data. For the H1 interface, AlphaFold produces predictions with high reliability scores that agree well with experimental data, while the H2 interface predictions appear inconsistent, preventing reliable binding modes. Despite these limitations, we experimentally assess the validity of Lana-1 and IE62 MED25 interface predictions. AlphaFold predictions also suggest the existence of a unique hydrophobic pocket for Arabidopsis MED25 ACID domain.

## Introduction

MED25 is a subunit specific to higher eukaryote Mediator complex, an essential component of the RNA polymerase II general transcriptional machinery (1). Many transcriptional activators have been reported to interact with human MED25 through its central activator interacting domain (ACID), also known as prostate tumor overexpressed (PTOV)/activator binding domain (ABD) (2, 3). This included the archetypal acidic transcriptional activation domain of the herpes virus activator VP16 (4–6), the Kaposi’s sarcoma-associated herpesvirus (KSHV) Lana-1 TAD (7), the varicella-zoster virus (VZV) major transactivator IE62 (8), the Respiratory Syncytial Virus Nonstructural Protein 1 (RSV NS1) (9–11) and the Ets related human transcription factors PEA3, ER81 and ERM (3, 12–14). In Arabidopsis, MED25 (AtMED25, also known as PFT1 (15)) also interacts with several transcriptional regulators and is considered as an integrator of multiple signaling pathways (16). Although dozens of partners have been described for Arabidopsis MED25 (17), there are no biophysical studies available besides those with AtDREB2a (18), VP16 (19) and AtMYC3 (20).

While the structure of human MED25 ACID domain has been solved by NMR by four different groups (2, 21–23), structures of MED25 ACID domain in complex with a target protein are still unknown. The human ACID domain contains a seven-stranded β-barrel flanked with three α helices and dynamic loops (Figure 1). Specifically, the human ACID domain coordinates its interactome using two largely hydrophobic interfaces named H1 et H2 and located on opposite faces of the β-barrel, surrounded by patches of positively charged residues. As defined by NMR chemical shift perturbation (CSP) and mutagenesis studies of ACID domain in complex with VP16 and ERM TADs (2, 13, 21, 22), the H1 binding site is delineated by strands β1-β3-β5, while the H2 binding site involves helix α1 and strands β6-β7. These ACID surfaces appear adequately designed to accommodate the specific patterns of bulky hydrophobic and negatively charged residues found in different acidic transactivation domains (2, 3, 10, 12, 13, 19, 21, 22, 24–26) (Figure 1).

**Figure 1:**
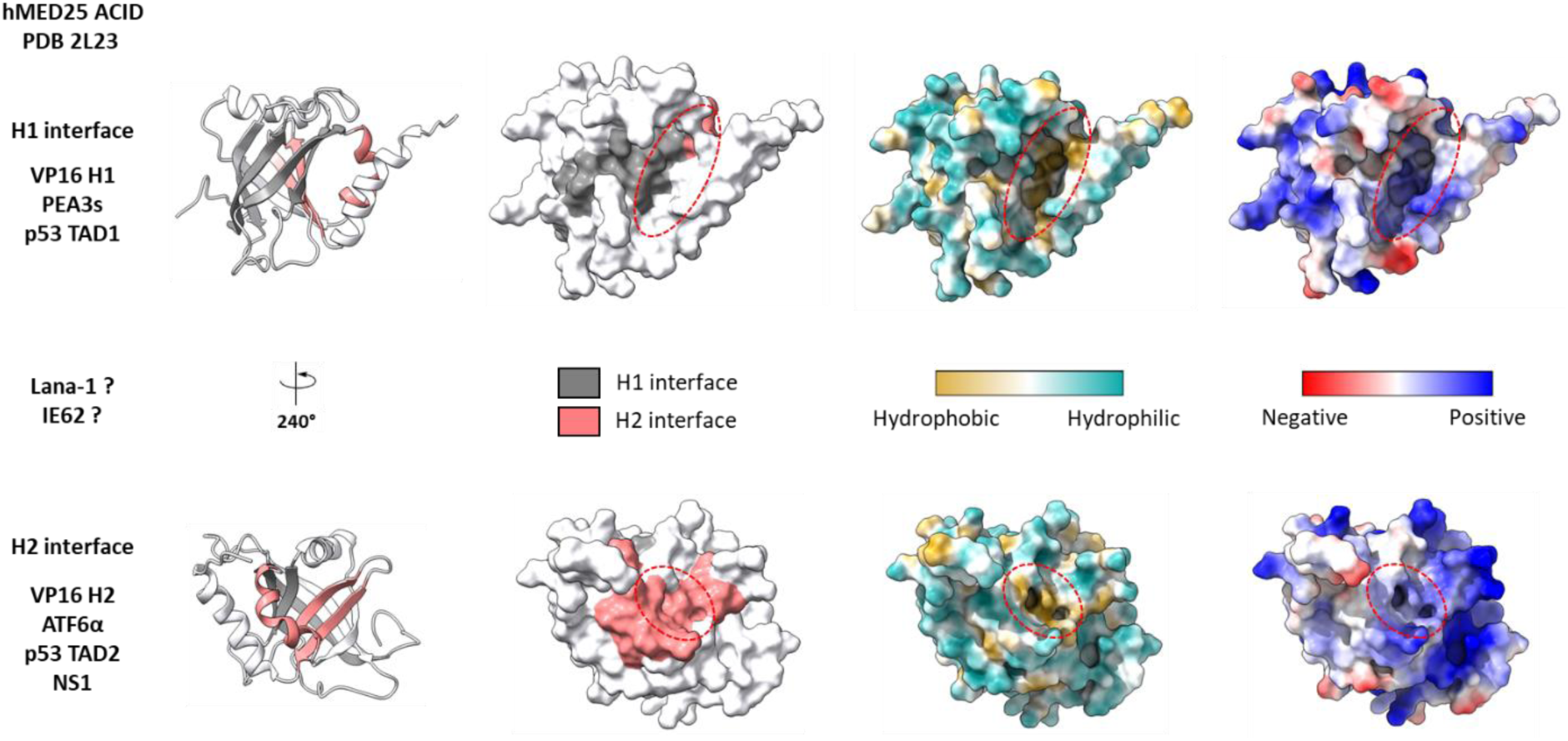
The human MED25 ACID domain. Cartoon and surface representation of human MED25 ACID domain (PDB 2L23 in white). The H1 binding site is formed mainly by strands β1-β3-β5 as defined by NMR chemical shift perturbation (21, 22) and is shown in gray. The H2 binding site is formed mainly by helix α1 and strands β6-β7 as defined by NMR chemical shift perturbation (2, 21, 22) and is shown in light coral. The various views of H1 and H2 are related by a 240° rotation along the y axis. PEA3s (ETV1/ER81, ETV4/PEA3 and ETV5/ERM) TADs, VP16 H1 subdomain and p53 TAD1 interact with the H1 face while VP16 H2 subdomain, ATF6α TAD, p53 TAD2 and the α3 helix of the respiratory syncytial virus NS1 protein interact with the H2 face. See main text for details and Supplementary Figure 1 for a summary of human MED25 ACID domain-TADs interactions. The binding interfaces of Lana-1 and IE62 are much less characterized as indicated by the question mark. The ACID domain surface is also colored by the molecular lipophilicity potential, where cyan denotes hydrophilic residues and gold denotes hydrophobic ones, and by electrostatic potential (blue and red indicate positively and negatively charged residues, respectively) as calculated with Coulombic surface coloring in ChimeraX (27). The two hydrophobic and positively charged pockets are indicated as a red dotted circle.

Numerous studies have helped to discriminate between MED25 partners according to the interface they contact. More specifically, the VP16 TAD subdomain H1, p53 subdomain TAD1 and PEA3s TADs directly interact with the H1 face of MED25 ACID (3, 12–14, 21, 22, 24, 25) while the VP16 TAD subdomain H2, p53 subdomain TAD2, ATF6α TAD and the C-terminal α3 helix of NS1 primarily bind to the MED25 H2 interface (2, 10, 21, 22, 24, 25). In contrast, the MED25 interaction surface and the minimal interaction domain for Lana-1 (7) and IE62 (8) remain largely unknow (Figure 1). A summary of the domain architecture of these MED25 interacting proteins with available experimental binding data is provided in Supplementary Figure 1.

In the absence of experimental structures of MED25-TADs complexes, computational modeling provides a valuable alternative. Machine learning (ML) approaches for *de novo* protein structure prediction such as neural networks AlphaFold2 (28) and RosettaFold (29) and large language model (LLM) OmegaFold (30) and ESMFold (31) have recently revolutionized structural biology by predicting highly accurate structures of proteins and their complexes.

In this study we report a systematic assessment of deep learning methods using the publicly available ColabFold version (32) to predict Human and Arabidopsis MED25 ACID domain-TADs interfaces and evaluate the accuracy of the models through comparison with published experimental data. We modelled 9 different human MED25 ACID-TAD complexes starting with the best-characterized experimentally (VP16, p53, ATF6α, ERM) and ending with the lesser-known ones (Lana-1 and IE62), enabling us to precisely predict and validate their interaction interfaces. Although AlphaFold was unable to discriminate between H1 and H2 interfaces, it remains a formidable hypothesis generator. The models correctly predict the minimal MED25 ACID interacting sequences of its binding partners that fold into α-helices in the vast majority of cases. We also reveal a new interaction surface unique to plants by predicting 3 different AtMED25 complexes (AtAP2/ERF, AtDREB2a and VP16). Finally, we compare the four main ML protein folding methods (AlphaFold, RoseTTAFold, OmegaFold and ESMFold) and discuss usability and limitations of their predictions.

## Materials and Methods

### Structural predictions

Protein sequences used in this study were extracted from the UniProt (33) database (Supplementary file 1). The predicted structures from AlphaFold, RoseTTAFold, OmegaFold and ESMFold were obtained (accessed on May and June 2023) from the python notebook available through the ColabFold interface (https://github.com/sokrypton/ColabFold) (32). An experimental notebook developed by Sergey Ovchinnikov (https://colab.research.google.com/github/sokrypton/ColabFold/blob/main/beta/omegafold_hacks.ipynb) was also used to incorporate Multiple Sequence Alignment (MSA) input into OmegaFold structure prediction (Figure 1). This notebook only supported monomeric and homo-oligomeric predictions and was therefore only used for the isolated human and Arabidopsis MED25 ACID domain (Figure 2). We also used a local installation of AlphaFold-multimer (version 2.3.0) or accessed directly from ChimeraX (version 1.6rc202303310103 (2023-03-31)) (27).

**Figure 2:**
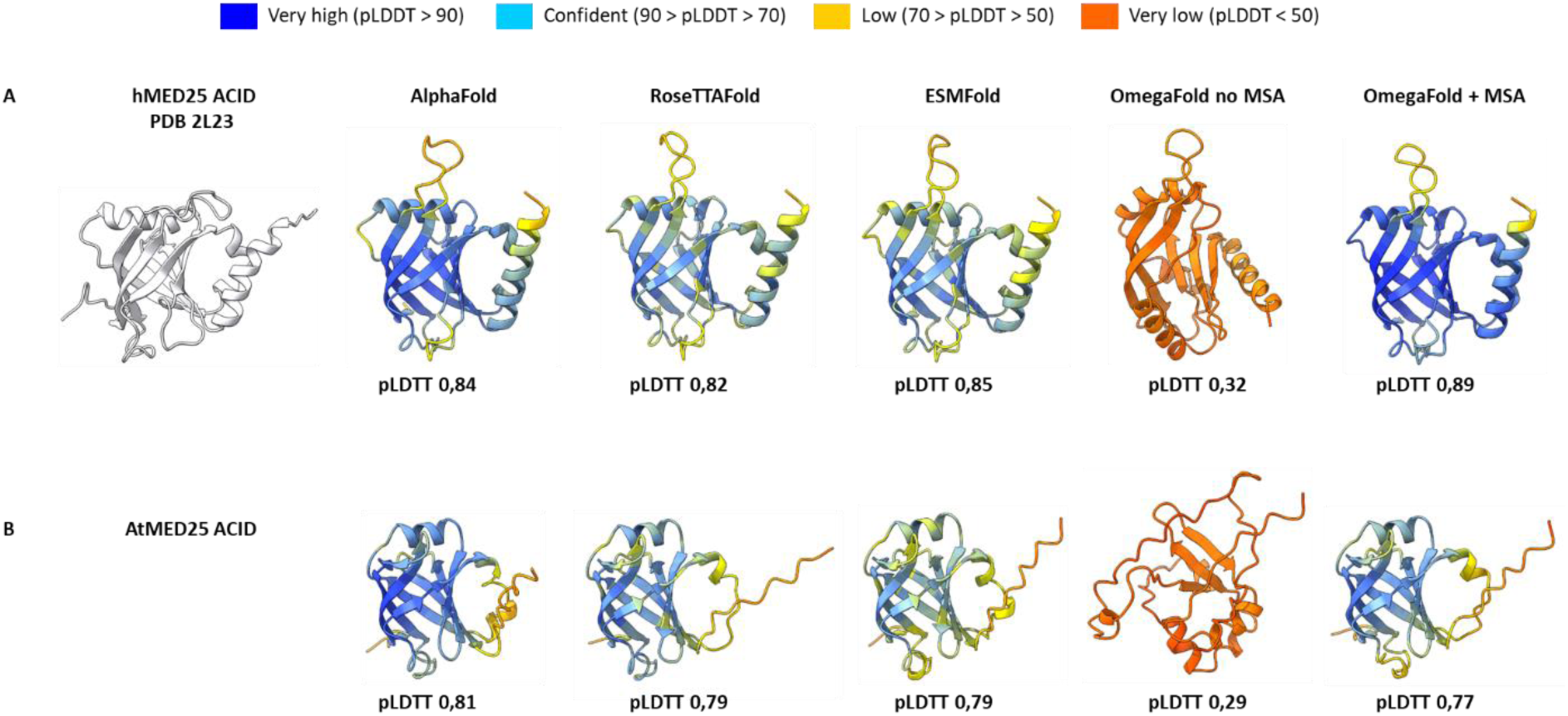
ML-based structural prediction of Human and Arabidopsis MED25 ACID domain. **(A)** Cartoon representation of human (PDB 2L23) and **(B)** Arabidopsis thaliana MED25 ACID domain and comparison of the initial structures obtained with machine learning protein folding methods AlphaFold, RoseTTAFold, ESMFold and OmegaFold. Each residue in human (hACID) and Arabidopsis (AtACID) MED25 ACID domain is color-coded based on the model confidence score, pLDDT.

For each prediction, five initial models were produced with default settings within multimer mode. To note, the python notebook does not use PDB templates (no template), thereby providing a totally naive structure prediction. Based on template modelling scores and predicted alignment error (PAE) values for each model, we then picked the top-ranked predicted structure of the complex as the final model for subsequent analysis. Visualization of the PAE for predicted complex structures was done with the webserver PAE Viewer (34) and figures were prepared using ChimeraX (27).

The protein-protein interactions modeled by AlphaFold-multimer distribute over the full pTM and ipTM range and multimer scores for the best ranked structures are shown in each figure and in Supplementary Figure 8. pTM, ipTM and mean pLDDT scores > 0.8 were considered highly accurate. For RoseTTAFold (29), mean interface predicted aligned error (pAE) < 5 indicated high confidence. The pLDDT (predicted local-distance difference test) is a confidence measure for the per-residue accuracy of the structure, the pTM (predicted template modeling score) is a metric for the similarity between protein structures (accuracy of prediction within each protein chain) and the ipTM (Interface pTM) scores interactions between residues of different chain to estimate the accuracy of interfaces (accuracy of the complex between chain) (28, 35, 36).

### Plasmids, protein expression and purification

The pET24d vector encoding human MED25 ACID domain (391–548)-6xHis has been previously described (2). BamHI/XhoI primers corresponding to Lana-1 (280–297) were annealed and cloned into pGEX 6P1 plasmid. For IE62 1-200, we applied the synthetic gene approach to synthesize (GenBank X04370.1) a codon-optimized pGEX 6P1 expression vector from GenScript. IE62 1-86 and 87-200 were amplified by PCR (template IE62 1-200), digested by BamHI/NotI and subcloned into the pGEX 6P1 vector. The bacterial expression vectors pGEX 6P1, pGEX 6P1 Lana-1 280-297, IE62 1-200, IE62 1-86 and IE62 87-200 (ampicillin) were co-transformed with pET24d hMED25 ACID-6xHis (kanamycin) in competent *E. coli* BL21 (DE3) strain and grown at 37°C in LB medium (150 ml) to an optical density of 0.8 at 600 nm and expression was induced with 0,1 mM IPTG for 18 h at 18°C. Cells were harvested by centrifugation and store frozen at -20°C. Pellets were resuspended in 12 ml cold lysis buffer (50 mM Tris-HCl pH 8,0, 100 mM NaCl, 10% glycerol and 0.1% NP-40 supplemented with 0.1 mg of lysozyme/ml and complete inhibitor cocktail (Roche)) and sonicated to disrupt the cells. The lysates were cleared by centrifugation at 10,000 × *g* at 4°C for 45 min to collect soluble proteins. The lysates were incubated with glutathione Sepharose 4B beads or TALON® Superflow^TM^ resin for 2h at 4°C and the beads were pelleted and washed three times with 10 volumes of lysis buffer. Bound proteins were analysed by mPAGE^TM^ 4-20% Bis-Tris gels (Merck) and Coomassie staining. Gels were scanned with Amersham ImageQuant 800.

## Results

### Benchmarking machine learning models for human and Arabidopsis MED25 ACID domain structure

We first assessed the accuracy of ML-based prediction algorithms AlphaFold (28), RoseTTAFold (29), OmegaFold (30) and ESMFold (31) to correctly predict the structure of hMED25 ACID domain alone. Unlike AlphaFold and RoseTTAFold, OmegaFold and ESMFold use large protein language model as a backbone and do not need a Multiple Sequence Alignment (MSA) step to generate prediction. The ColabFold version (32) was used with default settings (no template) to predict a set of five structural models of hMED25 ACID domain and the models with the highest overall pLDDT scores were picked (Figure 2A). Predictions showed an excellent agreement (with an average pLDDT score > 0.8 and a RMSD below 1Å) with the NMR structures (PDB 2L23, 2L6U, 2XNF and 2KY6) (2, 21–23) with the exception of OmegaFold, which only becomes accurate by incorporating MSA as additional input (Figure 2A).

The structure of AtMED25 ACID domain has not been experimentally determined and despite a low sequence identity with the human ACID domain, structure prediction indicates that the β-barrel architecture is topologically conserved (Figure 2B). AtMED25 ACID predicted β-strands β1-β4-β5-β7 and α helix H2 overlap quite well with their human counterpart with an RMSD value of 1.12 Å (Supplementary Figure 2). The major difference being a missing H3 helix in Arabidopsis in line with previous results (18). As with human ACID, OmegaFold prediction of AtACID appears only accurate with an additional MSA input (Figure 2B). Thus, ML-based structural prediction of MED25 ACID is in good agreement with NMR structures, indicating that the generated models are suitable for further analysis.

### Case study of human MED25 ACID domain structure prediction in complex with well-characterized interacting protein partners

We next tested if AlphaFold can predict the structure of human MED25 ACID domain in complex with partner proteins that were well characterized biochemically, but for which the structure of the complexes has not been experimentally resolved to date. We focused on the herpes virus activator VP16 (21, 22), the three Ets transcription factors ERM, PEA3 and ER81, (3, 12–14), p53 (25) and ATF6α (24, 37). These six MED25 partners represent an excellent starting point for our study since they are distributed across the entire spectrum of all possible interactions: bivalent MED25 interaction (H1 and H2 interfaces) for VP16 H1H2 and p53 TAD1 TAD2, MED25 H1 interface interaction for VP16 H1 subdomain, PEA3s TADs and p53 TAD1 and MED25 H2 interface interaction for VP16 H2 subdomain, p53 TAD2 and ATF6α TAD (Figure 1 and Supplementary Figure 1). In all cases, AlphaFold predicted an interface which is mediated by alpha helices formed by the acidic TADs, although with varying accuracy (Figure 3).

**Figure 3:**
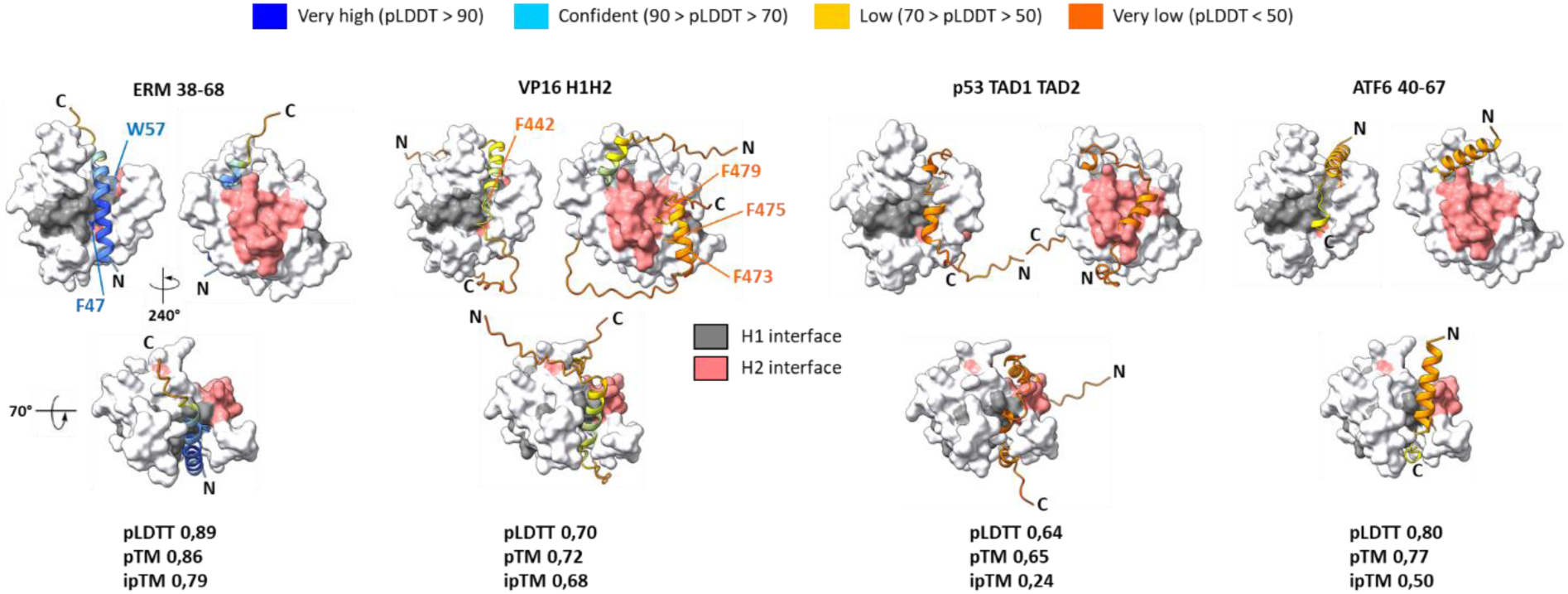
AlphaFold structural predictions of human MED25 ACID domain in complex with ERM, VP16, ATF6α and p53 transactivation domains. Surface representation of human MED25 ACID domain (white) with interfaces H1 (gray) and H2 (light coral) in complex with ERM, VP16, ATF6α and p53 transactivation domains (cartoon representation color-coded based on the model confidence score, pLDDT). N-terminal (N) and C-terminal (C) regions of the transactivation domains are indicated. Cartoon representation related by 240° rotation along the y axis and related by 70° rotation along the x axis. For each prediction, the pLDDT (predicted local-distance difference test), pTM (predicted template modeling score) and ipTM (Interface pTM) scores are indicated.

#### PEA3s

ERM (38–68) TAD has the highest ipTM (0,79) and pTM (0,86) scores supported by high pLDDT (0,89), indicating very high confidence in both the interface and in the prediction of the overall structure. The ERM TAD helical structure predicted by AlphaFold fitted into the MED25 H1 interface with residues Y47 (pLDDT 0,95) and W57 (pLDDT 0,90) that pointed toward the concave hydrophobic cavity (Figure 3), in agreement with experimental data (12, 13) (Supplementary Figure 1). Highly accurate predictions were also obtained with ER81/ETV1 (38–69) and PEA3/ETV4 (45–76) TADs (Supplementary Figure 3A and 3B). The sequence variations across the PEA3 subfamily TADs resulted in a slightly shorter helix for PEA3 which could explain the unique PEA3 MED25 engagement mode as compared to ERM and ER81 (3). In addition to the PEA3s TADs, MED25 ACID domain was also reported to interact with the α-helix H4 that is specific to ETV1/ETV4/ETV5 DNA binding domain (DBD) (14). Three distinct sites on MED25 bind the ETS DBD, two of which correspond roughly to the H1 and H2 interfaces (site 1 and site 2 in (14)). The MED25 ACID-PEA3 DBD (337–436) model has a low pTM score (0,57) and a very low iPTM score (0,18) (Supplementary Figure 3C), indicating poor confidence in the interface that roughly corresponds to the third site (site 3) identified in (14). Nevertheless, using full length PEA3 (1–484) as template (Supplementary Figure 4), AlphaFold correctly predicts two distinct binding surfaces for PEA3 on MED25 ACID.

#### VP16

AlphaFold successfully predicted the extended binding interface of VP16 with the H1 and H2 subdomains wrapping around the β-barrel and adopting a partially folded conformation when bound (Figure 3). Three separate regions with α-helical propensity are apparent spanning residues 428 to 441, 444 to 447 (VP16 H1 subdomain) and 469 to 482 (VP16 H2 subdomain) which included F442, F473, F475 and F479 that have been shown to be critical for VP16 transcriptional activity and MED25 recruitment (5, 21, 22, 38).

#### p53

In contrast to VP16, AlphaFold did not accurately predict the relative orientation of the MED25 ACID/p53 TAD1 TAD2 complex (Figure 3 and Supplementary Figure 1). The model correctly predicts two distinct binding surfaces but with the binding site for p53 TAD1 on the MED25 H2 interface and the binding site for p53 TAD2 on the H1 interface. The confidence scores are low (pLDTT 0,64, pTM 0,65, ipTM 0,24), suggesting that the MED25/p53 TAD model is unreliable.

#### ATF6α

We next modelled MED25/ATF6α TAD 40-67 complex that was reported as an H2 binding site specific partner (24) (Supplementary Figure 1). AlphaFold prediction suggested a preferential occupation of MED25 ACID by both a very short α-helix in the C-terminus of ATF6α (64-LDL-66) in the H1 interface and a second predicted long N-terminal α-helix (41-TDELQLEAANETYENNFDN-59) arranged in a near perpendicular fashion with the H1 α-helix of the ACID β-barrel (Figure 3). However, the global quality metrics are good but not very high for the ipTM score (pLDDT 0,8, pTM 0,77, ipTM 0,50), indicating limited reliability of the predicted binding interface. We also explored how predictions would be impacted by queries in which the putative bound ATF6α motif is embedded in larger fragments. Using ATF6α 1-80 (Supplementary Figure 5), 1-150 and 38-75 (data not shown) as templates (24, 37), AlphaFold predictions that were obtained are quite different from the initial ATF6α 40-67 prediction (Figure 3). In all cases, AlphaFold was unable to accurately predict the binding mode and to discriminate between the H1 and H2 ACID interfaces with many different predicted ATF6α short α-helixes fragments embedded in long disordered regions.

#### p53 TAD1, TAD2 and VP16 H1 and H2 subdomains

By comparing p53 TAD1, p53 TAD2, VP16 H1 and VP16 H2 subTADs boundaries, we also found variation in predictive performance among models (Figure 3 and Supplementary Figure 6). In all cases, higher accuracy models were generated but predicted interaction interfaces do not always overlap. While the VP16 H1 subdomain model is almost indistinguishable from the one with VP16 H1H2, the VP16 H2 subdomain prediction gives now 2 different solutions with very similar confidence scores, either VP16 H2 in the ACID H1 interface or in the H2 interface. For p53 TAD1 subdomain, AlphaFold now predicts the correct H1 interface but suggests the same solution for p53 TAD2 (Figure 3 and Supplementary Figure 6).

#### Comparison of machine-learning models

Likewise, comparing AlphaFold, RoseTTAFold and ESMFold, we found that ERM, VP16, p53 and ATF6α TADs models do not always converge (Figure 3, Supplementary Figures 5 and 7). For example, none of the tested ML protocols converged to an identical model for ATF6α (40–67) and ESMFold fitted incorrectly the ERM TAD into the H2 interface.

### Case study of respiratory syncytial virus NS1 protein

We next assessed the accuracy of AlphaFold, RoseTTAFold, and ESMFold at correctly predicting the structure of hMED25 ACID domain in complex with the respiratory syncytial virus NS1 protein. MED25 was recently identified as an NS1 interacting protein (9, 11) and NMR experiments indicate that the NS1 C-terminal α3 helix contacts directly the ACID H2 interface (10) (Supplementary Figure 1). RoseTTAFold and ESMFold predictions converged to an identical model for NS1 (118–139) bound to the H2 ACID domain interface that matches experimental data while AlphaFold fitted the NS1 C-terminal helix into the H1 interface (Figure 4). However, AlphaFold prediction has the highest pTM (0,80) score supported by high confidence pLDDT (0,82). This is intriguing in light of the NMR data, which showed that a peptide consisting of the sequence of the NS1 α3 helix primarily bound to MED25 ACID H2 with 17 µM affinity. However, the H1 interface was also detected as a secondary binding site with an apparent affinity of 500 µM (10).

**Figure 4:**
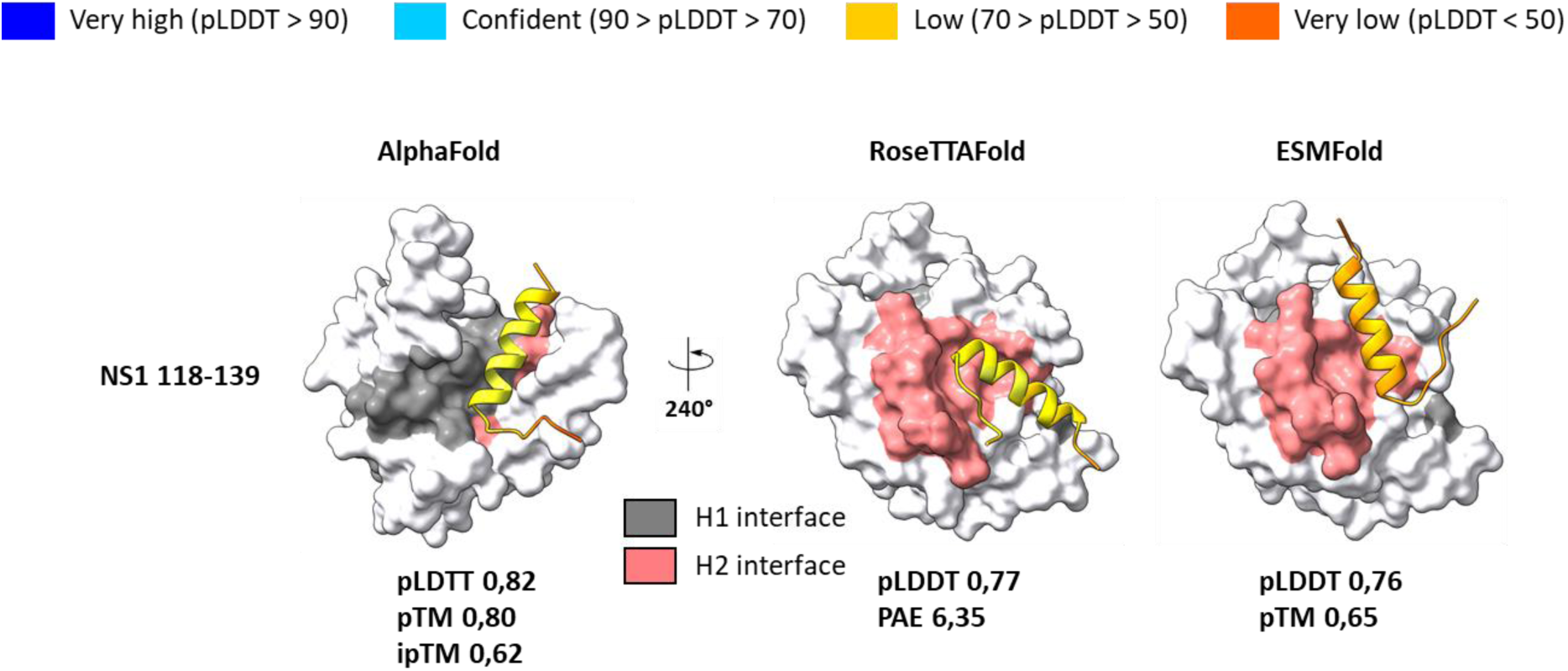
ML-based structural prediction of human MED25 ACID domain in complex with respiratory syncytial virus NS1 C-terminal α3 helix. Surface representation of human MED25 ACID domain (white) with the two opposite binding surfaces H1 (gray) and H2 (light coral) in complex with NS1 C-terminal helix (118–139) (cartoon representation color-coded based on the model confidence score, pLDDT). For each prediction, the pLDDT (predicted local-distance difference test), pTM (predicted template modeling score), PAE (predicted alignment error matrix) and ipTM (Interface pTM) scores are indicated.

### Case study of human MED25 ACID domain structure prediction in complex with poorly characterized interacting protein partners

We next used AlphaFold to map precisely the binding region of KSHV Lana-1 and VZV IE62 to MED25. Previous reports have shown an interaction between the N-terminal region of Lana-1 (1–340) (7) and the N-terminal region of IE62 (1-226 and 1-86) (8, 39) and MED25 ACID domain but the minimal domains of interaction have not yet been identified (Supplementary Figure 1).

#### Lana-1

The Lana-1 segment (1–340) is predicted to be almost entirely disordered by AlphaFold except for a long α-helix at the C-terminus (residues 279-306) fitted in the ACID H2 interface but the confidence scores are very low (pLDTT 0,43, pTM 0,39, ipTM 0,36) (Figure 5A). Delineating the input sequence into a shorter fragment (Lana-1 279-308) clearly increases the confidence metrics (pLDTT 0,82, pTM 0,79, ipTM 0,63) (Figure 5B). However, the predicted interaction interfaces are opposite from each other between Lana-1 (1–340) and Lana-1 (279–308), making the identification of the most plausible interface very challenging. Looking more closely, we noted that the ACID H1 interface is in fact occupied by Lana-1 (1–340) disordered region centered on residues Y235 and W238, probably rendering it inaccessible to the C-terminal predicted α-helix (Figure 5A). To experimentally verify the AlphaFold predictions and establish the contribution of this Lana-1 (279–308) putative interface, we co-expressed GST and GST Lana-1 (280–297) with human MED25 ACID domain in bacteria. Bacterial lysates were incubated with glutathione or cobalt beads, washed extensively and bound complexes were analysed by SDS-PAGE and Coomassie blue staining. As shown in Figure 5C, MED25 ACID was effectively and specifically pulled down by GST Lana-1 (280–297), thus validating the AlphaFold model.

**Figure 5:**
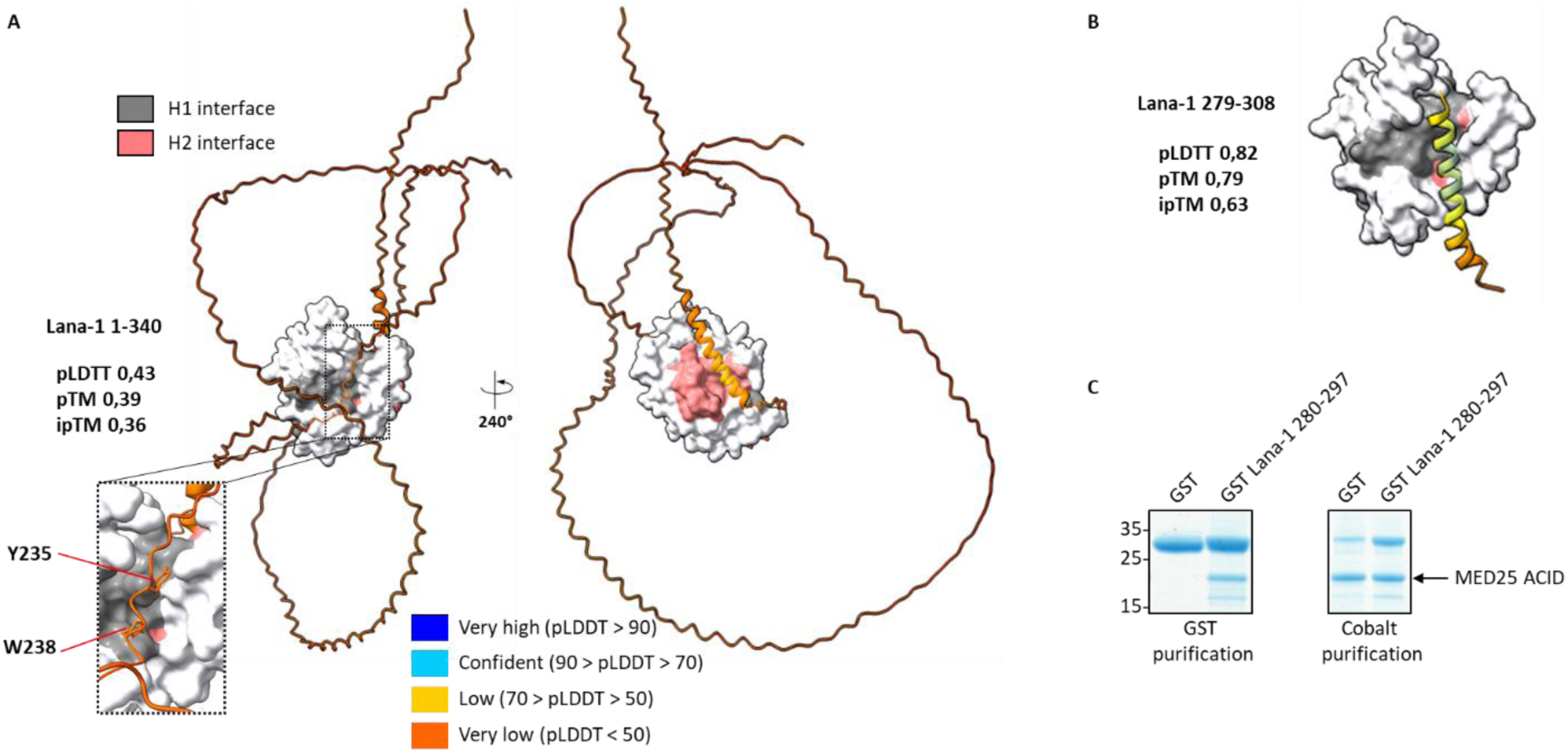
AlphaFold structural prediction of human MED25 ACID domain in complex with the Kaposi’s sarcoma-associated herpesvirus (KSHV) Lana-1 protein. **(A)** Surface representation of human MED25 ACID domain (white) with the two opposite binding surfaces H1 (gray) and H2 (light coral) in complex with **(A)** Lana-1 (1–340) and **(B)** Lana-1 (279–308) (cartoon representation color-coded based on the model confidence score, pLDDT). **(A)** Inset, close-up view of Lana-1 (1–340)-ACID H1 interface with Lana-1 residues Y235 and W238 highlighted. Each structure in H1 and H2 view related by 240° rotation along the y axis. For each prediction, the pLDDT (predicted local-distance difference test), pTM (predicted template modeling score) and ipTM (Interface pTM) scores are indicated. **(C)** Copurification of MED25 ACID with GST Lana-1 (280–297). GST Lana-1 (280–297) or GST were co-expressed with MED25 ACID-6xHis in bacteria and purified with glutathione Sepharose 4B beads (left) or cobalt resin (right). Fractions were analysed by SDS-PAGE and coomassie blue staining. As indicated with cobalt resin, MED25 ACID domain is expressed to similar level. Molecular weight marker (in kDa) is indicated.

#### IE62

A similar approach was used with the varicella-zoster virus (VZV) major transactivator IE62 (Figure 6). We first used IE62 (1–200) as template (Figure 6A) which roughly corresponds to the (1–226) domain initially described as interacting with MED25 (39). AlphaFold predicted an extended binding interface reminiscent of VP16 H1H2 and p53 TAD1 TAD2 with two partially folded α-helix wrapping around MED25 ACID β-barrel, while the remaining part of IE62 (1–200) appeared mostly unstructured (Figure 6A). The predicted IE62 N-terminal α-helix (residues 17-43) fitted in the ACID H1 interface (Figure 6A) is in good agreement with previous experimental data that identified a potential extended α-helix involving residues 19-35 of IE62 TAD (8). The second predicted IE62 α-helix (residues 106-117) bound to the H2 interface (Figure 6A) has never been considered. The confidence scores are low (pLDTT 0,53, pTM 0,53, ipTM 0,68) but given the fact that disordered regions lack proper structure, the model quality of those regions will be always low, hence lowering the overall model quality (40). Using IE62 1-86, 99-200 (Figure 6A), 17-41 and 106-120 (Figure 6B) as templates, AlphaFold predictions that were obtained are quite different from the initial IE62 1-200 prediction. In all cases, AlphaFold predicted an H2 interface binding mode. To establish the relative contribution of these IE62 putative interfaces, we co-expressed GST, GST IE62 (1–200), (1–86) and (87–200) with human MED25 ACID domain in bacteria. Bacterial lysates were incubated with glutathione or cobalt beads, washed extensively and bound complexes were analysed by SDS-PAGE and Coomassie blue staining. As shown in Figure 6C, MED25 ACID was effectively and specifically pulled down only by GST IE62 (1–200) and (1–86), indicating that the IE62 99-200 and IE62 106-120 MED25 ACID models are incorrectly classified as interacting pairs by AlphaFold.

**Figure 6:**
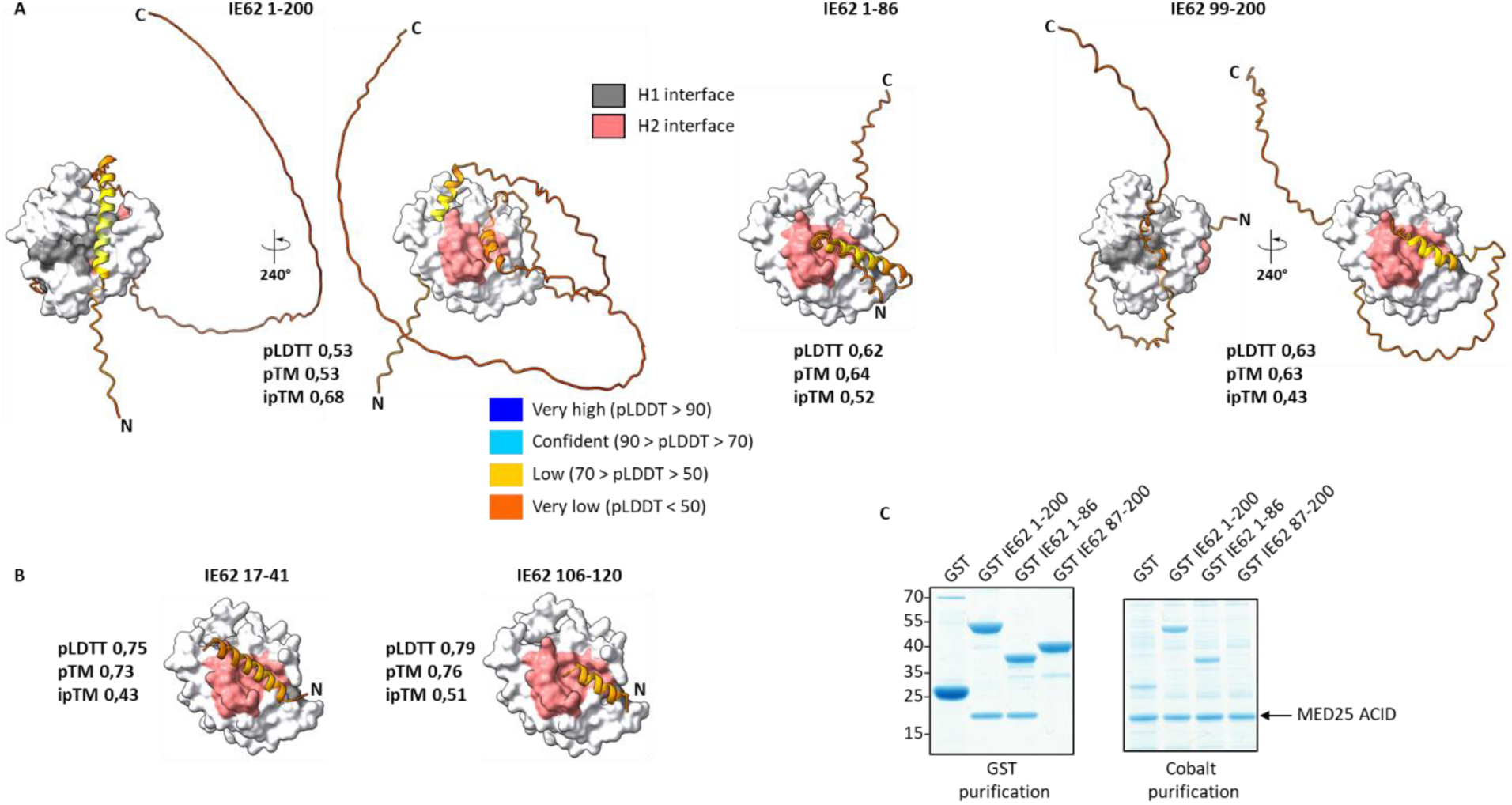
AlphaFold structural prediction of human MED25 ACID domain in complex with the varicella-zoster virus (VZV) major transactivator IE62. Surface representation of human MED25 ACID domain (white) with the two opposite binding surfaces H1 (gray) and H2 (light coral) in complex with various fragments of IE62 (cartoon representation color-coded based on the model confidence score, pLDDT). **(A)** IE62 (1–200), (1–86) and (99–200) predictions. **(B)** IE62 (17–41) and (106–120) predictions. N-terminal (N) and C-terminal (C) regions of the transactivation domains are indicated. Each structure in H1 and H2 view related by 240° rotation along the y axis. For each prediction, the pLDDT (predicted local-distance difference test), pTM (predicted template modeling score) and ipTM (Interface pTM) scores are indicated. **(C)** IE62-MED25 ACID interaction by GST pull-down assay. GST IE62 (1–200), (1–86) and (87–200) or GST were co-expressed with MED25 ACID-6xHis in bacteria and purified with glutathione Sepharose 4B beads (left) or cobalt resin (right). Fractions were analysed by SDS-PAGE and coomassie blue staining. As indicated with cobalt resin, MED25 ACID domain is expressed to similar level. Molecular weight marker (in kDa) is indicated.

### Case study of Arabidopsis MED25 ACID domain structure prediction in complex with interacting protein partners

We finally tested if AlphaFold can predict the structure of Arabidopsis MED25 ACID domain in complex with selected interacting proteins. We focused on the Ethylene Response factors (AP2/ERF) family (16), the transcription factor AtDREB2a (18, 41) and the herpes virus activator VP16 (19).

In all cases, AlphaFold predicted an interface which is mediated predominantly by α helices, although with varying accuracy (Figure 7A). AtERF98 (117–139) TAD has the highest ipTM (0,82) and pTM (0,86) scores supported by high pLDDT (0,88), indicating very high confidence in both the interface and in the prediction of the overall structure. A C-terminal AtERF98 helical structure predicted by AlphaFold (DKVLEELLDSEERK) occupied a hydrophobic pocket defined by the AtMED25 ACID H2 α-helix and strands β1-β2 loop located at the top of the β-barrel core (Figures 7A and 7C and Supplementary Figure 2). This interface differs greatly from those determined for human MED25, likely due to the fact that there is no predicted H3 α-helix in AtMED25 and therefore no equivalent of the human H1 interface (Figure 1 and Supplementary Figure 2). The AlphaFold model suggested a second potential interface (Supplementary Figure 9) between AtACID and AtERF98 with a short N-terminal AtERF98 β-strand (FEFEY) that associates with an extra AtACID β-strand (β8 VVFKP) not predicted in AtMED25 ACID structure alone (Figure 1 and Supplementary Figure 2), reminiscent of a two-stranded β zipper (42). Nevertheless, the predicted AtMED25 ACID/AtERF98 complex matched experimental data closely with the conserved EDLL motif previously implicated in transcriptional activation and interaction with AtMED25 (16, 43) adopting a partially folded conformation when bound (Figure 7A and Supplementary Figure 9). In contrast, confidence scores are very low for VP16 H1H2 TAD, VP16 H1 and H2 subdomains (Figure 7A), making the identification of the most plausible interface very challenging.

**Figure 7:**
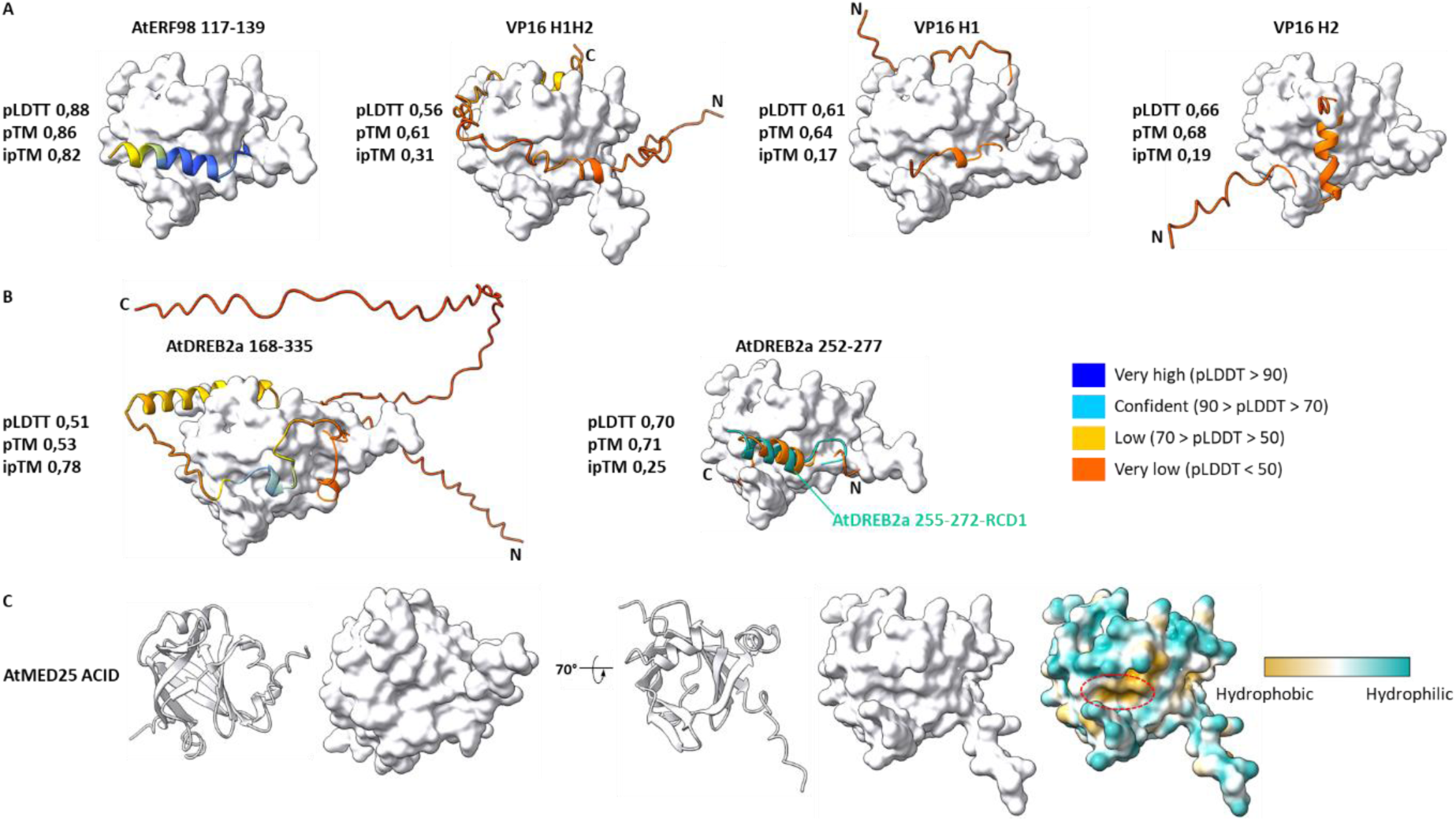
AlphaFold structural prediction of Arabidopsis thaliana MED25 ACID domain in complex with selected partners. **(A)** Surface representation of AtMED25 ACID domain (white) in top view in complex with AtERF98 (117–139), VP16 H1H2, VP16 H1 and VP16 H2 and **(B)** with AtDREB2a (168–335) and AtDREB2a (252–277) (cartoon representation color-coded based on the model confidence score, pLDDT). N-terminal (N) and C-terminal (C) regions of the transactivation domains are indicated. For each prediction, the pLDDT (predicted local-distance difference test), pTM (predicted template modeling score) and ipTM (Interface pTM) scores are indicated. The AtMED25 ACID-AtDREB2a (252–277) AlphaFold model was superimposed with the HADDOCK model of the RCD1-RST-DREB2A (255–272) complex. RCD1-RST was omitted for simplicity and AtDREB2a (255–272) α-helix is colored in light sea green. **(C)** Cartoon and surface representation of AtMED25 ACID domain as in Figure 1. Each structure in front and top view related by 70° rotation along the x axis. AtACID domain surface is also colored by the molecular lipophilicity potential, where cyan denotes hydrophilic residues and gold denotes hydrophobic ones with ChimeraX (27). The top hydrophobic pocket is indicated as a red dotted circle.

We next modelled AtMED25/AtDREB2a (168–335) complex (Figure 7B) that was reported to interact with a K_d_ in micromolar range by SPR (18) and ITC (19). This C-terminal AtDREB2a fragment comprised both the BD (168–253) and AD (254–335) domains but only the extended (168–335) fragment interacted with AtMED25 ACID with significant affinity (18, 19). AlphaFold was unable to accurately predict the binding mode with many different predicted AtDREB2a (168–335) α-helices embedded in long disordered regions bound to multiple AtMED25 ACID surfaces (Figure 7B). Interestingly, VP16 and AtDREB2a that both bind AtMED25, display very low sequence identity apart from a short positively charged stretch of amino acids in DREB2a (252-277 HLDSSDMFDVDELLRDLNGDDVFAGL) which lies exactly at the junction between the (168–253) and (254–335) domains (18, 19). Delineating the input sequence into this shorter 252-277 fragment clearly increases pLDTT and pTM confidence metrics (Figure 7B). Importantly, the 3 different complexes (ERF98, DREB2a and VP16) predicted here occupied the same AtMED25 hydrophobic pocket (Figure 7). Interestingly, AtDREB2a has also been reported to interact with the plant Radical-Induced Cell Death1 (RCD1) protein through the same C-terminal region that partially adopt an α-helical conformation upon association with RCD1 (44, 45). When our AlphaFold AtMED25-DREB2a (252–277) model was superimposed with the HADDOCK model of AtRCD1 in complex with DREB2a (255–272) (44), the two AtDREB2a α-helices aligns well (Figure 7B). Taken together, our results predicted a unique hydrophobic pocket located at the top edge of the AtMED25 β-barrel core, not conserved with hMED25 ACID, that is targeted by AtAP2/ERF family, AtDREB2a and VP16 (Figure 7C). The observation that AtDREB2a and VP16 interacted with overlapping regions of AtMED25 ACID (19) and that the MED25 ACID- and RCD1-binding regions of DREB2a partly embraced (44, 45) supported our AlphaFold modeled structures.

## Discussion

In this paper, we compared machine-learning algorithms for the structure prediction of several acidic transactivation domains in complex with Mediator complex subunit MED25. This case study is particularly well suited for testing the current limits of artificial intelligence, as it combines a number of challenges: (i) The MED25 ACID domain is structurally unique among activator binding domains (21) with its seven-stranded β barrel core flanked by 3 α-helices, and consequently might be underrepresented in the training set of AlphaFold, (ii) TADs are intrinsically disordered, poorly conserved and fold into multiple orientation and conformation when bound (46–49), (iii) TADs acquired structure might vary with context such as the nature of the binding partner (50) and (iv) AlphaFold models have limited conformational variance for now. To address these limitations, different approaches are currently under development to increase the structural heterogeneity of the predicted ensemble (51).

An overview of the MED25 ACID/TADs structural models produced here shows that the large majority of them form between a defined putative short α-helix residing in the intrinsically disorder TAD and the structured MED25 ACID β core domain. However, all the protein– protein interactions modeled by AlphaFold distribute over the full pTM and ipTM range, with only a subpopulation of highly confident predictions with ipTM and pTM > 0.80 (Supplementary Figure 8). This suggests that some complexes can’t be correctly predicted regardless of the input conditions, most notably all the partners who are supposed to interact with the MED25 H2 interface. Several hypotheses can explain the low confidence scores associated with the folded ML model segments. First, the observed differences in modeling performance can be due to the intrinsic characteristics of each AI model in particular with the language model ESMFold (31) and OmegaFold (30) that do not require multiple sequence alignment and templates. Second, structure predictions may be unreliable due to the lack of representation in the training set or an insufficient amount of homologous sequences to validate the predicted contacts. In particular, it has been shown recently (52) that the percentage of the predicted disordered region in the sequences of unassigned domains had an inverse correlation coefficient with model quality for AlphaFold and RoseTTAFold models, indicating that model quality are lower because the sequences might be disordered. Third, as previously observed with the yeast acidic transcription activator Gcn4-MED15 complex (46, 53, 54) and human MED25-PEA3s (3, 13, 14, 26), TADs likely bind MED25 ACID in multiple conformations and orientations that prevents accurate prediction of their correct binding mode. It is also possible that AlphaFold training procedure may have been biased by overrepresented experimental structures in the PDB. On closer inspection, we noticed an almost systematic presence of an aromatic residue in the MED25 interacting partners that may constrain the machine learning algorithms to propose a ‘preferred’ solution in interaction with the H1 interface (Supplementary Figure 10). Superimposition of the MED25 ACID/TADs complexes shows that in the majority of cases, tyrosine or tryptophan residues have their side chains pointing in the same direction with ERM (38–68) F47 residue (Supplementary Figure 10). Nevertheless, the signature of TAD folding is there and thus, given that some proteins such as Lana-1, IE62 and the majority of Arabidopsis MED25 protein partners remain largely uncharacterized, AlphaFold predictions constitute interesting working hypotheses. As prediction of complexes is, on average, less accurate than that of folds of individual proteins, it is particularly important to always verify interface predictions experimentally as we have done here with IE62 and Lana-1. Our results also underscore the interest of fragment-based searching to identify interaction motifs when one of the protein partners is predicted highly disordered. In case where the binding region is unknown, delineating the interaction region into fragments of decreasing size may increase the overall success rate in agreement with a recent analysis (55).

The observation that VP16 H1 subdomain, PEA3s TADs and NS1 C-terminal α3-helix bind the same MED25 ACID H1 interface despite low sequence identity suggests that conformational plasticity within the 7-strands β-barrel core domain could play a role in mediating partner recognition. Transient kinetics experiments and molecular dynamics simulations clearly support that MED25 ACID-TAD complexes are highly dynamic (24). In particular, strands β1-β2 loop (residues 409-424) and α-helix H3 (residues 529-543) flanking the H1 binding interface are the most dynamic region of the ACID domain and show the greatest stabilization upon binding (24). AlphaFold predictions of ERM TAD (Figure 3) and NS1 C-terminal α3-helix (Figure 4) in complex with MED25 ACID support the structural model of binding seen from molecular dynamics. Superimposition of VP16 H2-, ERM (38–68)- and NS1 (118–139)-bound MED25 models suggest opening of the H1 hydrophobic groove where ACID α-helix H3 shifts away to accommodate the different relative positions of the TAD α-helical conformation (Supplementary Figure 11). Our results also support the recently proposed acidic exposure model for TAD function in which acidic residues and intrinsic disorder keep hydrophobic motifs exposed to solvent where they are available to bind coactivators (56–59).

In this work, we also predicted the structure of Arabidopsis MED25 ACID domain in complex with three different interacting proteins. The predicted shared binding interface located at the top of the β-barrel core, led us to hypothesize that hMED25 and AtMED25 ACID are structurally conserved (Figure 1) but that AtMED25-TADs interactions are specific to plants (Figure 7). During the preparation of our manuscript, an independent biophysical analysis of AtMED25 ACID domain in complex with AtDREB2a became available (60). Our results with AtDREB2a (252–277) are in good agreement with the NMR, ITC, MD and AlphaFold experiments performed in this study with a longer AtDREB2a fragment (234–276). In particular, the authors identified two AtMED25 ACID binding motifs in AtDREB2a (234–276) (60), one of which corresponds to the one we have just highlighted in this study. Additional AlphaFold Arabidopsis MED25 ACID domain/TADs models to guide experimental studies will help elucidate how sequences divergence between orthologs may play a role in dictating molecular recognition.

## Acknowledgements

We thank Christina Sizun for critically reading the manuscript and Sergey Ovchinnikov for the OmegaFold_hacks experimental notebook.

## Notes

### Competing Interest Statement

The authors have declared no competing interest.

