## Supplementary Figures for "Assessment of machine-learning predictions for MED25 ACID domain interactions with transactivation domains"

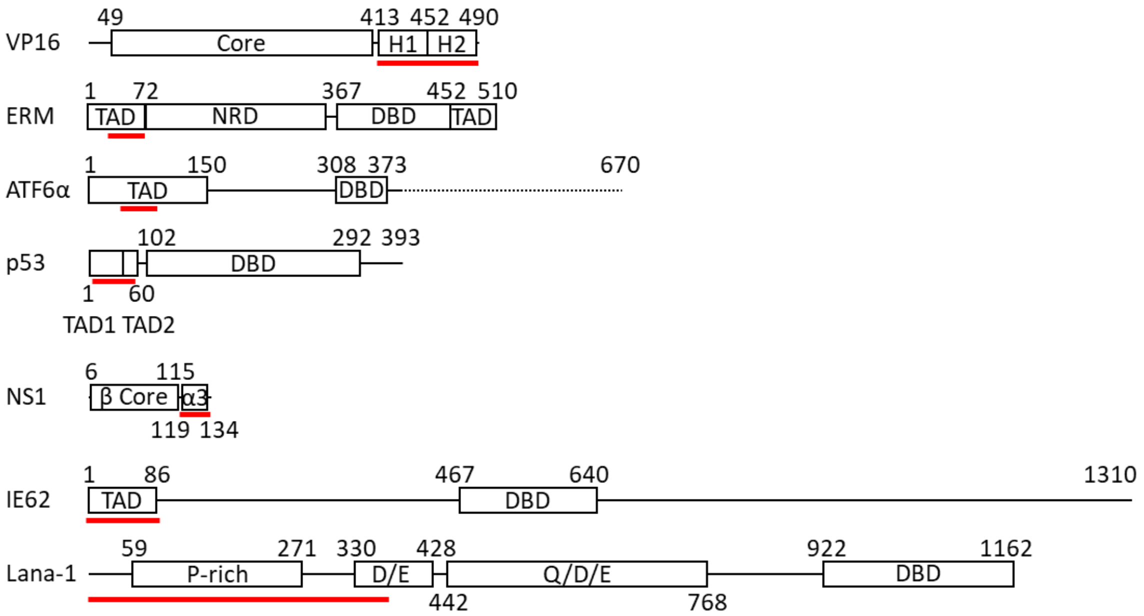


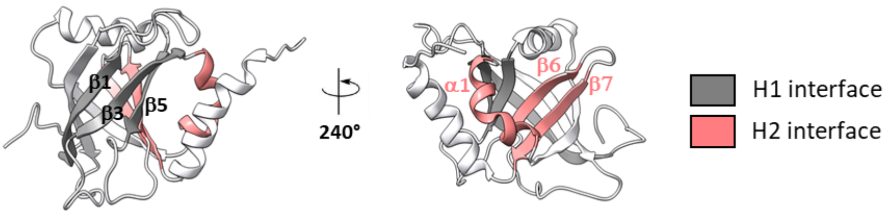


| Protein | Residues | MED25 Interface | Techniques | References |
|---|---|---|---|---|
| VP16 H1 H2 | 411-490 | H1 & H2 | Pull-down, CSP, ITC, FP, stopped-flow, MD | (3-5, 12-13, 19, 21-22) |
| VP16 H1 | 411-452 | H1 | Pull-down, CSP, ITC, FP, stopped-flow, MD | (3-5, 12-13, 19, 21-22) |
| VP16 H2 | 453-490 | H2 | Pull-down, CSP, ITC, FP, stopped-flow, MD | (2-6, 19, 21-22, 24) |
| ERM | 38-68 | H1 | Pull-down, CSP, ITC, FP, stopped-flow, MD, Co-IP | (3, 12-14, 24, 26) |
| ER81 | 38-69 | H1 | Pull-down, CSP, stopped-flow | (14, 24) |
| PEA3 | 45-76 | H1 | Pull-down, CSP, stopped-flow | (14, 24) |
|  | 337-436 | multiple | CSP, BLI, ChIP | (14) |
| ATF6α | 40-66 | H2 | Pull-down, CSP, stopped-flow, MD | (24, 37) |
| p53 TAD1 TAD2 | 1-73 | H1 & H2 | CSP, ITC | (25) |
| p53 TAD1 | 15-29 | H1 | CSP | (25) |
| p53 TAD2 | 39-57 | H2 | CSP | (25) |
| NS1 | 118-139 | H2 | Y2H, CSP, ITC | (9-11) |
| IE62 | 1-86 | unknown | Pull-down, Co-IP | (8, 39) |
| Lana-1 | 1-340 | unknown | Pull-down | (7) |

**<u>Supplementary Figure 1:</u> Human MED25 ACID domain interactions with transactivation domains.** Various functionally characterized domains of VP16, ERM, ATF6α, p53, NS1, IE62 and Lana-1 used in this study and their location are shown as boxes. The numbers indicate amino acid positions. The minimal interaction domain for each of the proteins that interacted with MED25 ACID is shown as a red line. **VP16**: Core (Oct-1/HCF binding domain), H1 and H2 TAD subdomains. **ERM**: N-terminal TAD and C-terminal TAD, NRD (negative regulatory domain), DBD (ETS DNA-binding domain). **ATF6α**: TAD and DBD (DNA-binding domain). ATF6α is a membrane-bound transcription factor that activates genes in the endoplasmic reticulum (ER) stress response. When unfolded proteins accumulate in the ER, ATF6α is cleaved to release its cytoplasmic domain (shown as a dotted line). The N-terminal fragment translocates to the nucleus where it functions as a transcription factor. **p53**: TAD1, TAD2 and DBD. **NS1:** β core and C-terminal α3 helix. **IE62**: TAD and DBD. **Lana-1**: Proline-rich region (P-rich), DBD, acidic repeat regions (D/E) and (Q/D/E). A cartoon representation of human MED25 ACID domain is shown as in Figure 1 with strands β1-β3-β5 (H1 binding site) and helix α1 and strands β6-β7 (H2 binding site) highlighted. **Table**: Summary of VP16, PEA3s, ATF6α, p53, NS1, IE62 and Lana-1 interactions with human MED25 ACID domain characterized by different techniques. CSP: NMR Chemical shift perturbation, ITC: Isothermal titration calorimetry, FP: Fluorescence polarization, MD: Molecular dynamics, BLI: Bio-layer Interferometry, ChIP: Chromatin Immunoprecipitation, Co-Ip: Co-Immunoprecipitation, Y2H: Yeast two hybrid.

A

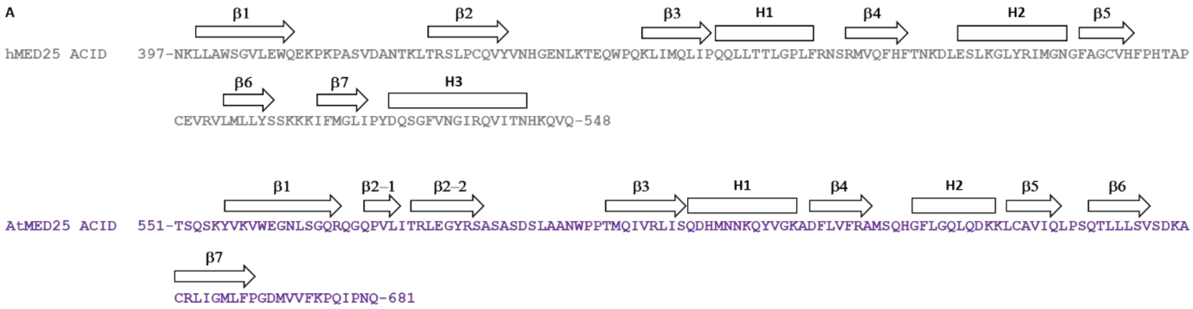


B

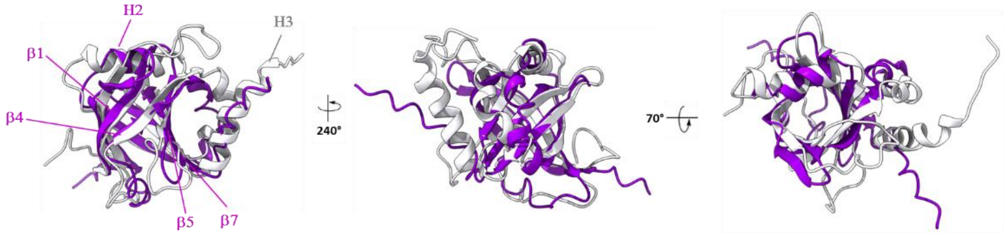


**<u>Supplementary Figure 2:</u> Structural similarity between human and Arabidopsis MED25 ACID domain. (A)** Primary and secondary structure of human (397-548 in gray) and Arabidopsis (551-681 in dark violet) MED25 ACID domain. Secondary elements (α-helices and β-strands) determined by NMR for human MED25 and predicted by AlphaFold for Arabidopsis MED25 are indicated above the sequences. **(B)** Superimposition of the experimentally determined structure of human MED25 ACID domain (397-548 in white, PDB 2L23) with the structure of Arabidopsis MED25 ACID domain as predicted by AlphaFold (551-681 in dark violet). Cartoon representation related by 240° rotation along the y axis and related by 70° rotation along the x axis. The root-mean-square deviation (RMSD) of mainchain atoms is 1.12 Å. Arabidopsis MED25 ACID β-strands β1-β4-β5-β7 and α helix H2 overlap well with human MED25 ACID and lacks the C-terminal helix H3.

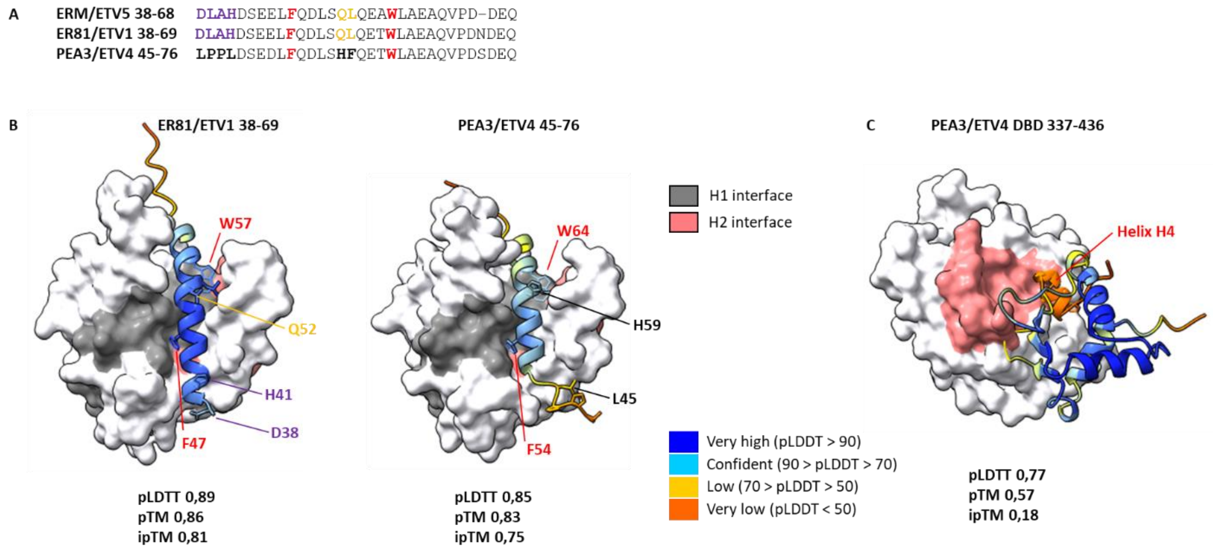


<u>**Supplementary Figure 3:**</u> **AlphaFold structural prediction of human MED25 ACID domain in complex with the PEA3 Ets members family. (A)** Alignment of ERM, ER81 and PEA3 transactivation domains. Residues that are variable between ERM, ER81 and PEA3 and the two conserved aromatic residues (F47 and W57 in ERM and ER81, F54 and W64 in PEA3) that are involved in the binding of MED25 are indicated. **(B)** Surface representation of MED25 ACID domain (white) with the H1 binding surface (gray) in complex with the minimal ER81/ETV1 and PEA3/ETV4 transactivation domains in cartoon representation color-coded based on the model confidence score, pLDDT. Residues that are variable between ERM, ER81 and PEA3 and the two conserved aromatic residues are indicated and color coded as in (A). **(C)** Surface representation of MED25 ACID domain (white) with the H2 binding surface (light coral) in complex with the DNA binding domain (DBD) of PEA3 (337-436) in cartoon representation color-coded based on the model confidence score, pLDDT. The additional α helix H4 of the DNA binding domain that is specific to the PEA3 subfamily and interacts with MED25 ACID domain is indicated. The predicted interface corresponds roughly to the unique site 3 described in (14). For each prediction, the pLDDT (predicted local-distance difference test), pTM (predicted template modeling score) and ipTM (Interface pTM) scores are indicated.

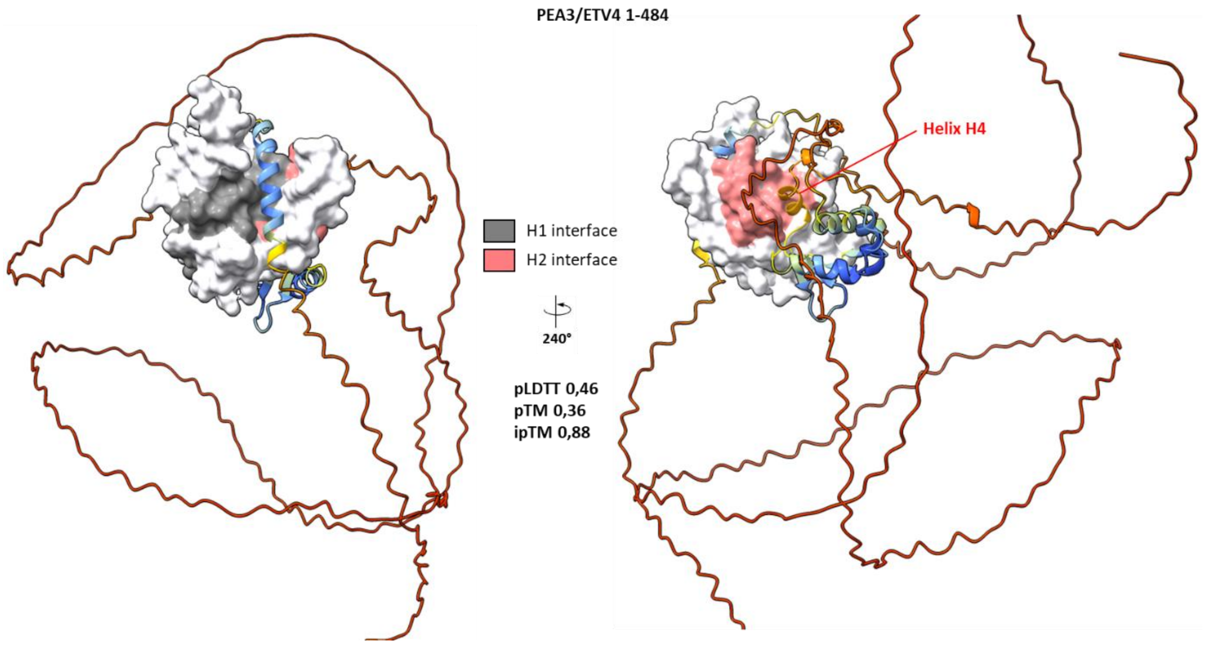


**<u>Supplementary Figure 4:</u> AlphaFold structural prediction of human MED25 ACID domain in complex with full length PEA3 (1-484).** Surface representation of MED25 ACID domain (white) with the H1 binding surface (gray) and the H2 binding surface (light coral) in complex with full length PEA3/ETV4 (1-484) in cartoon representation color-coded based on the model confidence score, pLDDT. Cartoon representation related by 240° rotation along the y axis. The additional α helix H4 of the DNA binding domain that is specific to the PEA3 subfamily and interacts with MED25 ACID domain is indicated as in supplementary Figure 3. The pLDDT (predicted local-distance difference test), pTM (predicted template modeling score) and ipTM (Interface pTM) scores are indicated.

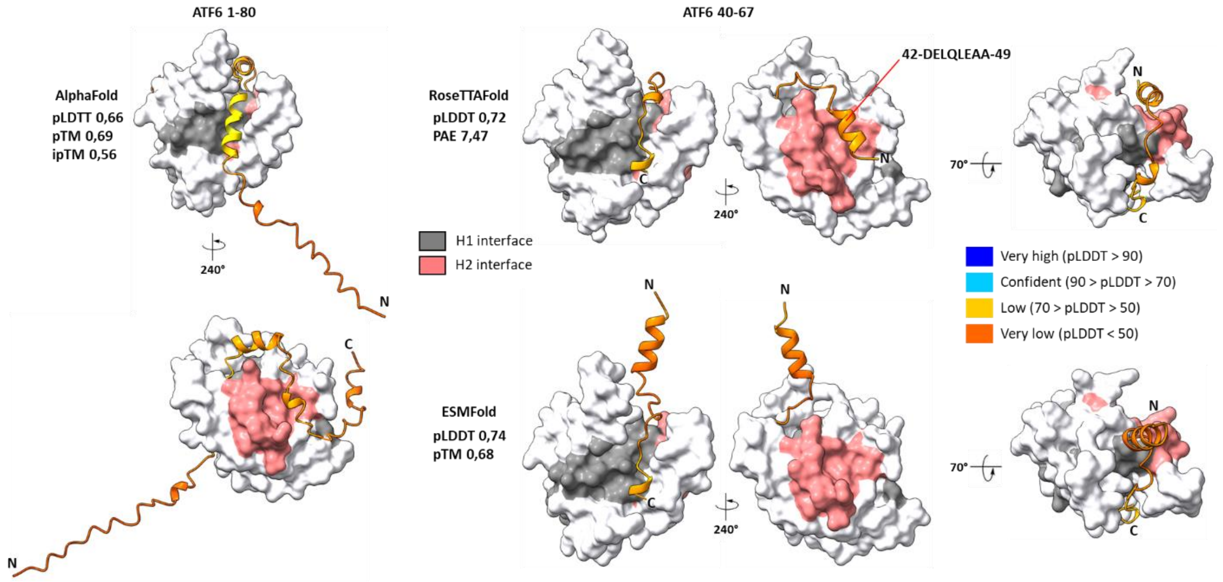


**<u>Supplementary Figure 5:</u> ML-based structural prediction of human MED25 ACID domain in complex with ATF6α transactivation domain.** Surface representation of MED25 ACID domain (white) with the H1 binding surface (gray) and the H2 binding surface (light coral) in complex with ATF6α (1-80) and (40-67) transactivation domains in cartoon representation color-coded based on the model confidence score, pLDDT. AlphaFold prediction (left) for ATF6α (1-80) and RoseTTAFold or ESMFold predictions (right) for ATF6α (40-67) are indicated. Cartoon representation related by 240° rotation along the y axis and related by 70° rotation along the x axis. For each prediction, the pLDDT (predicted local-distance difference test), pTM (predicted template modeling score), PAE (predicted alignment error matrix) and ipTM (Interface pTM) scores are indicated.

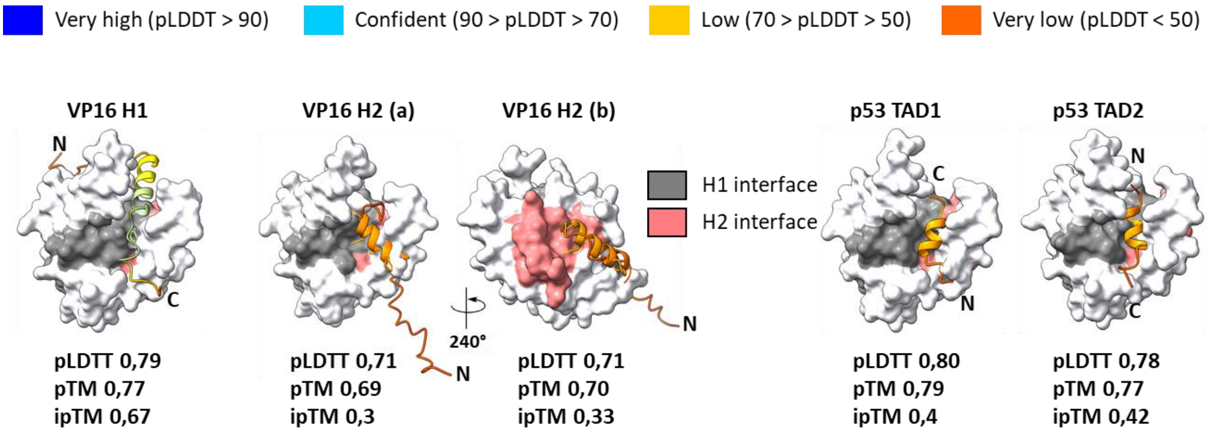


**Supplementary Figure 6: AlphaFold structural prediction of human MED25 ACID domain in complex with VP16 H1, VP16 H2, p53 TAD1 and p53 TAD2 transactivation subdomains.** Surface representation of MED25 ACID domain (white) with the H1 binding surface (gray) and the H2 binding surface (light coral) in complex with VP16 H1, VP16 H2, p53 TAD1 and p53 TAD2 subdomains in cartoon representation color-coded based on the model confidence score, pLDDT. Cartoon representation related by 240° rotation along the y axis. For each prediction, the pLDDT (predicted local-distance difference test), pTM (predicted template modeling score) and ipTM (Interface pTM) scores are indicated.

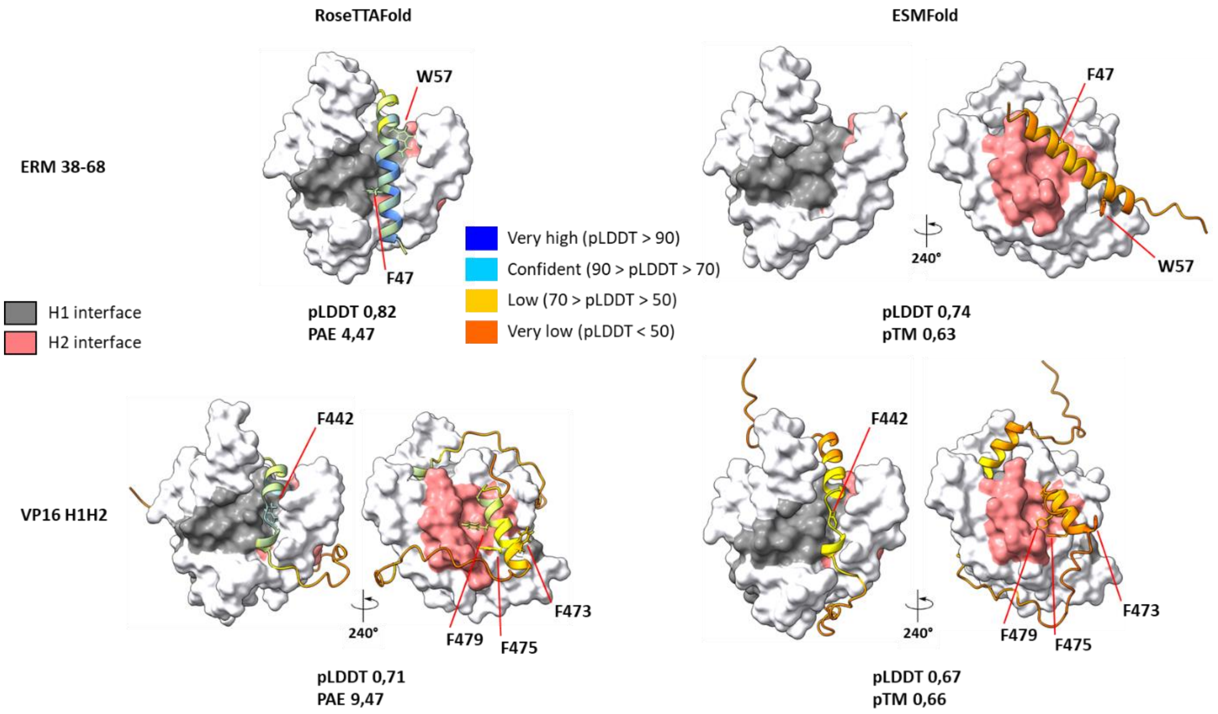


<u>**Supplementary Figure 7:**</u> **RoseTTAFold and ESMFold structural prediction of human MED25 ACID domain in complex with ERM 38-68 and VP16 H1H2 transactivation domains.** Surface representation of MED25 ACID domain (white) with the H1 binding surface (gray) and the H2 binding surface (light coral) in complex with ERM 38-68 and VP16 H1H2 transactivation domains in cartoon representation color-coded based on the model confidence score, pLDDT. (Left) RoseTTAFold prediction. (Right) ESMFold prediction. ERM (38-68) F47 and W57 residues, VP16 F442 (VP16 H1), F473, F475 and F479 (VP16 H2) residues that are involved in the binding of MED25 are indicated. Cartoon representation related by 240° rotation along the y axis. For each prediction, the pLDDT (predicted local-distance difference test), pTM (predicted template modeling score), PAE (predicted alignment error matrix) and ipTM (Interface pTM) scores are indicated.

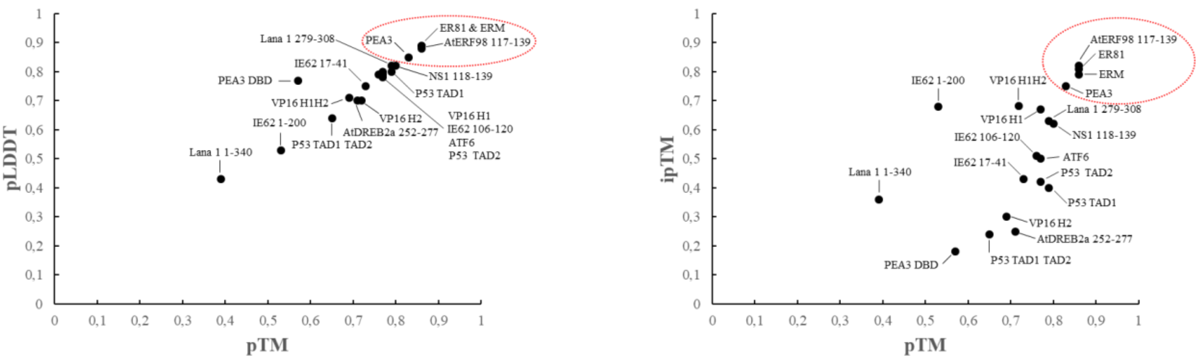


**<u>Supplementary Figure 8:</u> Structural prediction of human MED25 ACID domain in complex with transactivation domains with AlphaFold multimer.** (Left) Scatter plot displaying the relationship between pLDDT (y-axis) and pTM (x-axis) and (Right) Scatter plot displaying the relationship between ipTM (y-axis) and pTM (x-axis). The 19 protein–protein interactions (PPIs) modeled by AlphaFold-Multimer distribute over the full pTM and ipTM range, with a subpopulation of highly confident predictions with pLDDT, ipTM and pTM > 0.80 (indicated as a red dotted circle).

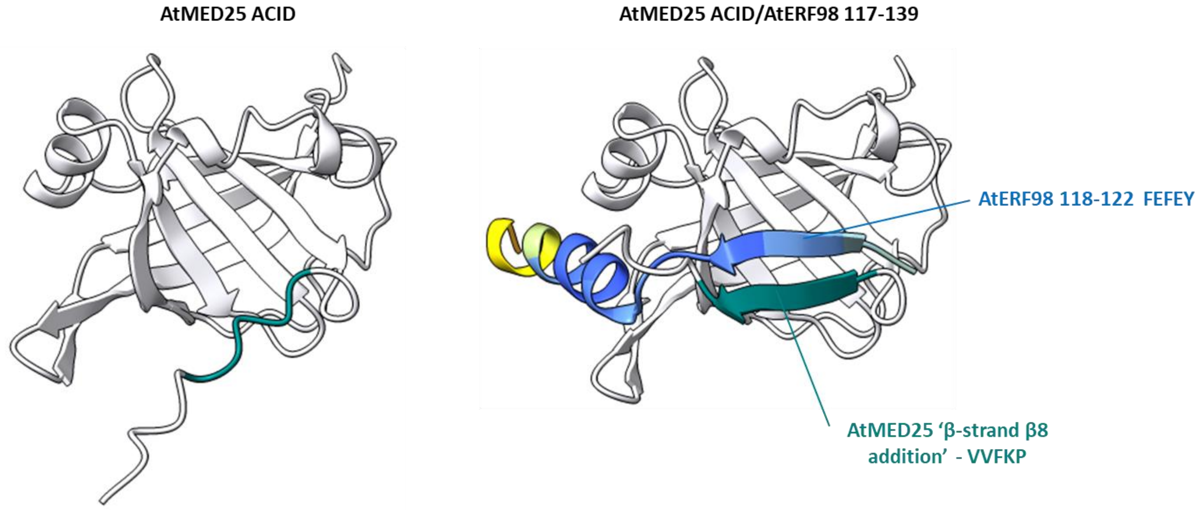


<u>**Supplementary Figure 9:**</u> **The AtMED25 ACID/AtERF98 (117-139) interface.** Cartoon representation of AtMED25 ACID domain (white) in top view related by 70° rotation along the y axis with respect to Figure 7. (Left) structure of AtMED25 ACID alone and (right) in complex with AtERF98 (117-139). The C-terminal region of AtMED25 ACID (VVFKP) that is predicted unstructured alone and predicted to form an extra β-strand (β8) in complex with AtERF98 (117-139) is indicated and colored in teal. The predicted N-terminal β-strand of AtERF98 (118-122 FEFEY) is also indicated.

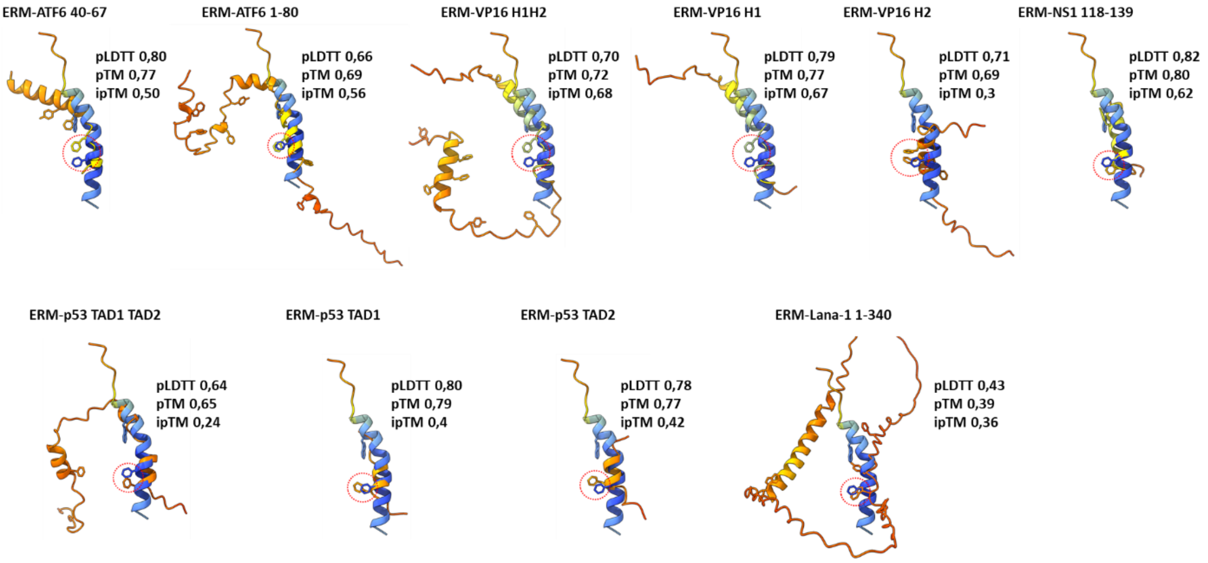


**Supplementary Figure 10: The ACID H1 binding pocket paradigm.** Comparison of the predicted ERM (38-68) TAD structure with all the other TADs in cartoon representation color-coded based on the model confidence, pLDDT. Each structure in H1 view were superimposed upon the NMR structure (PDB 2L23). The MED25 ACID domain was omitted for simplicity. For each prediction, the pLDDT (predicted local-distance difference test), pTM (predicted template modeling score) and ipTM (Interface pTM) scores are indicated. The Y and W hydrophobic residues of all the TADs are indicated and those that aligned with ERM F47 are indicated as a red dotted circle.

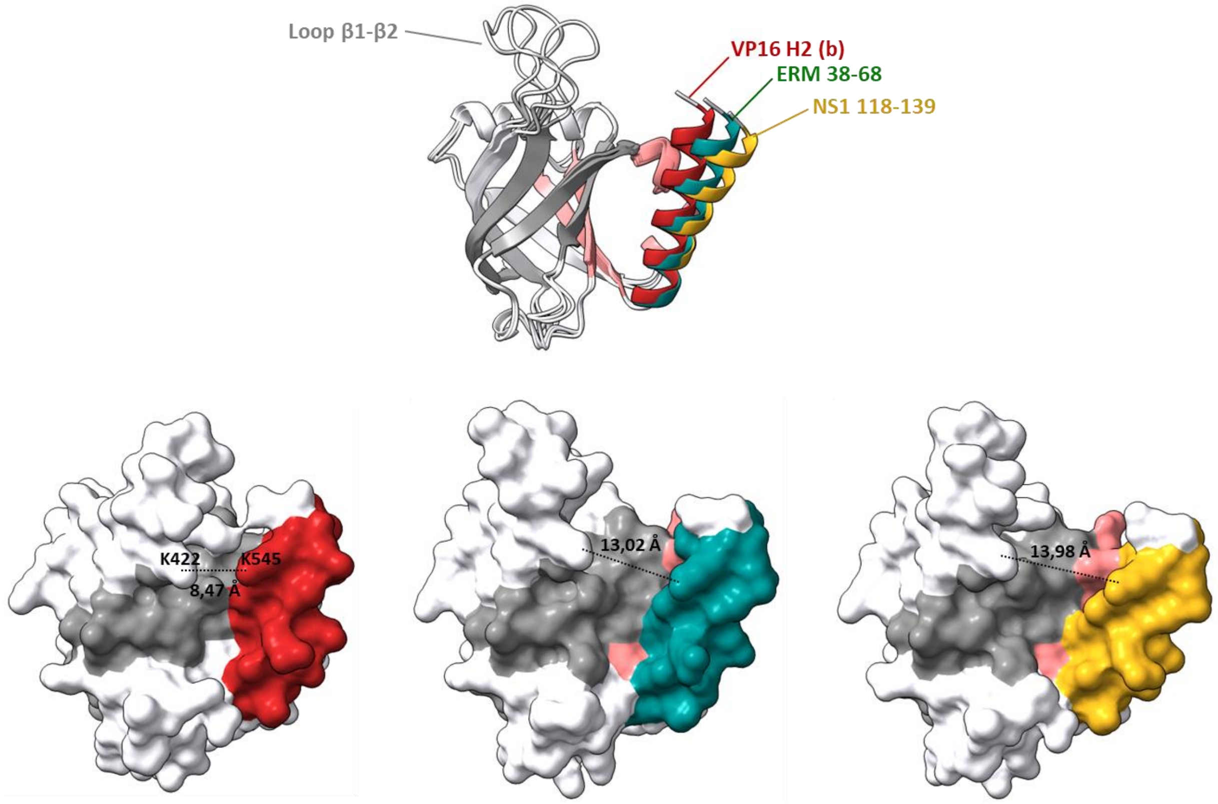


**<u>Supplementary Figure 11:</u> The ACID H1 binding pocket dynamic.** Cartoon and surface representation of MED25 ACID domain (white) with the H1 binding surface (gray) and the H2 binding surface (light coral). AlphaFold predictions of VP16 H2 (b) subdomain, ERM 38-68 TAD and NS1 C-terminal α3 helix in complex with MED25 ACID were superimposed upon the NMR structure (PDB 2L23, not shown). VP16 H2, ERM 38-68 and NS1 (118-139) were omitted for simplicity. The differences in ACID α3 helix location are highlighted by color corresponding to the TAD that is bound (crimson VP16 H2 (b), teal ERM 38-68 and goldenrod NS1 118-139). The flexible β1-β2 loop (409-424) of MED25 ACID domain is indicated. Average distance between residues K422 and K545 is indicated with black dashed line and values were measured with ChimeraX.
