## Supplementary file for "Assessment of machine-learning predictions for MED25 ACID domain interactions with transactivation domains"

**Supplementary file 1.** Protein sequences and UniProtKB accession numbers used in this study. TADs residues predicted to be composed of alpha helix by AlphaFold are highlighted in yellow.

**Q71SY5 hMED25 ACID NMR 2L23 397-548**

NKLLAWSGVLEWQEKPKPASVDANTKLTRSLPCQVYVNHGENLKTEQWPQKLIMQLIPQQLL
TTLGPLFRNSRMVQFHFTNKDLESLKGLYRIMGNGFAGCVHFPHTAPCEVRVLMLLYSSKKK
IFMGLIPYDQSGFVNGIRQVITNHKQVQ

**Q7XYY2 aMED25/PFT1 ACID domain 551-681**

TSQSKYVKVWEGNLSGQRQGQPVLITRLEGYRSASASDSLAANWPPTMQIVRLISQDHMNNK
QYVGKADFLVFRAMSQHGFLGQLQDKKLCAVIQLPSQTLLLSVSDKACRLIGMLFPGDMVVF
KPQIPNQ

**P41161 hERM/ETV5 38-68**

DLAHDSEELFQDLSQLQEAWLAEAQVPDDEQ

**P43268 hPEA3/ETV4 45-76**

LPPLDSEDLFQDLSHFQETWLAEAQVPDSDEQ

**P43268 hPEA3/ETV4 337-436**

RRGALQLWQFLVALLDDPTNAHFIAWTGRGMEFKLIEPEEVARLWGIQKNRPAMNYDKLSRS
LRYYYEKGIMQKVAGERYVYKFVCEPEALFSLAFPDNQ

**P43268 hPEA3/ETV4 1-484**

MERRMKAGYLDQQVPYTFSSKSPGNGSLREALIGPLGKLMDPGSLPPLDSEDLFQDLSHFQE
TWLAEAQVPDSDEQFVPDFHSENLAFHSPTTRIKKEPQSPRTDPALSCSRKPPLPYHHGEQC
LYSSAYDPPRQIAIKSPAPGALGQSPLQPFPRAEQRNFLRSSGTSQPHPGHGYLGEHSSVFQ
QPLDICHSFTSQGGGREPLPAPYQHQLSEPCPPYPQQSFKQEYHDPLYEQAGQPAVDQGGVN
GHRYPGAGVVIKQEQTDFAYDSDVTGCASMYLHTEGFSGPSPGDGAMGYGYEKPLRPFPDDV
CVVPEKFEGDIKQEGVGAFREGPPYQRRGALQLWQFLVALLDDPTNAHFIAWTGRGMEFKLI
EPEEVARLWGIQKNRPAMNYDKLSRSLRYYYEKGIMQKVAGERYVYKFVCEPEALFSLAFPD
NQRPALKAEFDRPVSEEDTVPLSHLDESPAYLPELAGPAQPFGPKGGYSY

**P50549 hER81/ETV1 38-69**

DLAHDSEELFQDLSQLQETWLAEAQVPDNDEQ

**P06492 HHV11 VP16 H1 413-452**

APPTDVSLGDELHLDGEDVAMAHADALDDFDLDMLGDGDS

**P06492 HHV11 VP16 H2 453-490**

PGPGFTPHDSAPYGALDMADFEFEQMFTDALGIDEYGG

**P06492 HHV11 VP16 H1H2 413-490**

APPTDVSLGDELHLDGEDVAMAHADALDDFDLDMLGDGDSPGPGFTPHDSAPYGALDMADFE
FEQMFTDALGIDEYGG

**P18850 hATF6 40-67**

DTDELQLEAANETYENNFDNLDFDLDLM

**P18850 hATF6 1-80**

MGEPAGVAGTMESPFSPGLFHRLDEDWDSALFAELGYFTDTDELQLEAANETYENNFDNLDF
DLDLMPWESDIWDINNQI

**P04637 hp53 TAD1 TAD2 1-60**

MEEPQSDPSVEPPLSQETFSDLWKLLPENNVLSPLPSQAMDDLMLSPDDIEQWFTEDPGP

**P04637 hp53 TAD1 15-29**

SQETFSDLWKLLPEN

**P04637 hp53 TAD2 46-60**

SPDDIEQWFTEDPGP

**P09310 HHV3 IE62/ICP4 1-200**

MDTPPMQRSTPQRAGSPDTLELMDLLDAAAAAAEHRARVVTSSQPDDLLFGENGVMVGREHE
IVSIPSVSGLQPEPRTEDVGEELTQDDYVCEDGQDLMGSPVIPLAEVFHTRFSEAGAREPTG
ADRSLETVSLGTKLARSPKPPMNDGETGRGTTPPFPQAFSPVSPASPVGDAAGNDQREDQRS
IPRQTTRGNSPGLP

**P09310 HHV3 IE62/ICP4 1-86**

MDTPPMQRSTPQRAGSPDTLELMDLLDAAAAAAEHRARVVTSSQPDDLLFGENGVMVGREHE
IVSIPSVSGLQPEPRTEDVGEELT

**P09310 HHV3 IE62/ICP4 99-200**

MGSPVIPLAEVFHTRFSEAGAREPTGADRSLETVSLGTKLARSPKPPMNDGETGRGTTPPFP
QAFSPVSPASPVGDAAGNDQREDQRSIPRQTTRGNSPGLP

**P09310 HHV3 IE62/ICP4 17-41**

PDTLELMDLLDAAAAAAEHRARVVT

**P09310 HHV3 IE62/ICP4 106-120**

LAEVFHTRFSEAGAR

**P0DOE9 HRSVA NS1 118-139**

SDSTMTNYMNQLSELLGFDLNP

**Q9QR71 HHV8P Lana-1 1-340**

MAPPGMRLRSGRSTGAPLTRGSCRKRNRSPERCDLGDDLHLQPRRKHVADSVDGRECGPHTL
PIPGSPTVFTSGLPAFVSSPTLPVAPIPSPAPATPLPPPALLPPVTTSSSPIPPSHPVSPGT
TDTHSPSPALPPTQSPESSQRPPLSSPTGRPDSSTPMRPPPSQQTTPPHSPTTPPPEPPSKS
SPDSLAPSTLRSLRKRRLSSPQGPSTLNPICQSPPVSPPRCDFANRSVYPPWATESPIYVGS
SSDGDTPPRQPPTSPISIGSSSPSEGSWGDDTAMLVLLAEIAEEASKNEKECSENNQAGEDN
GDNEISKESQVDKDDNDNKDDEEEQETDEE

**Q9QR71 HHV8P Lana-1 279-308**

DTAMLVLLAEIAEEASKNEKECSENNQAGE

**Q9LTC5 aERF98/TDR1 117-139**

VFEFEYLDDKVLEELLDSEERKR

**O82132 aDREB2a 168-335**

DCESKPFSGGVEPMYCLENGAEEMKRGVKADKHWLSEFEHNYWSDILKEKEKQKEQGIVETC
QQQQQDSLSVADYGWPNDVDQSHLDSSDMFDVDELLRDLNGDDVFAGLNQDRYPGNSVANGS
YRPESQQSGFDPLQSLNYGIPPFQLEGKDGNGFFDDLSYLDLEN

**O82132 aDREB2a 252-277**

HLDSSDMFDVDELLRDLNGDDVFAGL
